# Phytoassisted synthesis of biogenic silver nanoparticles using *Vernonia amygdalina* leaf extract: characterization, antibacterial, anti-inflammatory, and acute toxicity profile

**DOI:** 10.1101/2024.10.31.621348

**Authors:** Fonye Nyuyfoni Gildas, Belle Ebanda Kedi Philippe, Ntoumba Agnes Antoinette, Nkana Blondeau Parfait, Djuidje Annie Guilaine, Tako Djimefo Alex Kevin, Evouna Madeleine Ines Danielle, Nanga Chick Christian, Chimi Tchatchouang Geordamie, Kang Eha-Kang Costly, Paboudam Gbambie Awawou, Eya’ane Meva Francois

## Abstract

Increasing antimicrobial resistance and pathological consequences from long-term anti-inflammatory drug use necessitate the urgent discovery and rational design of new, effective antimicrobial and anti-inflammatory agents. This study reports the antibacterial, anti-inflammatory, and acute toxicity profile of *Vernonia amygdalina* leaf extract-mediated silver nanoparticles (VA-AgNPs). VA-AgNPs were synthesized by mixing Ag^+^ ions with an aqueous leaf extract from *Vernonia amygdalina* and characterized using UV-Visible (UV-Vis) spectroscopy, powder X-ray diffraction (PXRD), and scanning electron microscopy (SEM). Antibacterial activity against *Escherichia coli* strains, carrageenan-induced rat paw edema, and oral acute toxicity assays were performed. The change in color from light brown to dark brown indicated the formation of VA-AgNPs, which was further confirmed by the surface plasmon resonance peaks between 400 and 450 nm in the UV-Vis spectrum. Stability studies showed increased VA-AgNPs formation at basic pH and with higher reactant quantities. The X-ray pattern showed nanocrystalline particles of Ag and AgCl with mean sizes of 13 and 18 nm, respectively. The SEM images depict aggregates of spherical shapes. The synthesized VA-AgNPs inhibited *Escherichia coli* growth with a minimum inhibition concentration of 0.125 mg/mL and rat paw edema with a maximum inhibition percentage of 96% at a dose of 0.4 mg/kg body weight. Oral administration of VA-AgNPs was not associated with any toxicity, supporting their use in the development of new effective antibacterial and anti-inflammatory drugs.

## INTRODUCTION

Recent advances in science and technology have enabled the development of nanomaterials with at least one dimension in the nanoscale range, which exhibit unique or enhanced properties compared to their bulk counterparts. [1]. Although nanomaterials have existed for centuries, their diverse applications in biomedicine, cosmetics, pharmaceutics, and energy have recently attracted growing scientific interest [2]. Metal nanoparticles are highly attractive because of their tunable physical and chemical properties, including surface plasmon excitation, which makes them ideal for applications in drug delivery. [3]. Bacterial infections pose a global public health burden, and rising antimicrobial resistance threatens to cause severe consequences. To address this, numerous studies have explored the development of antibacterial nanoparticles as novel therapeutic agents, highlighting their advantages over conventional methods and leveraging diverse natural resources for the synthesis of bionanomaterials. [4]. Colloidal silver nanoparticles have been used as antimicrobial agents, wound dressing materials, bone and tooth cements, and water purifiers [5]. Recent efforts have focused on developing green chemistry methods for the synthesis of silver nanoparticles with the advantage of using natural products and avoiding toxic reducing agents, organic solvents, and wasteful purifications with high residual cytotoxicity [6–8].

*Vernonia amygdalina*, commonly known as bitter leaf, is a shrub that grows up to 2-5 meters high in African countries, including Cameroon, Zimbabwe, and Nigeria [9]. It belongs to the Asteraceae family, with a petiolate leaf of approximately 6 mm in diameter and an elliptical shape [10]. The plant is traditionally used to treat amoebic dysentery and gastrointestinal disorders and has antimicrobial and antiparasitic activities [9]. The biologically active compounds of *Vernonia amygdalina* include saponins, coumarins, flavonoids, lignans, alkaloids, and xanthones [11]. This study highlights the potent antibacterial activity of optimized *Vernonia amygdalina*-derived silver nanoparticles against *E. coli*, alongside their anti-inflammatory properties and acute toxicity profile to establish a preliminary safety profile providing insights on their therapeutic potential.

## MATERIALS AND METHODS

### Bacteria

*Escherichia coli* strains were isolated from urine samples in the bacteriology unit of the General Hospital laboratory, Yaoundé, Cameroon.

### Plant Extract

Fresh bitter leaves (*Vernonia amygdalina*) were harvested on July 14, 2020, at Logpom, Littoral region, Cameroon (Figure 1) and authenticated at the National Herbarium of Cameroon, in comparison with a voucher specimen previously deposited (N°32875 HNC). *V. amygdalina* leaves were cleaned with tap water, then deionized water, to remove dust and particles. The aqueous extract of *V. amygdalina* was prepared by boiling 25 g of finely cut leaves in 250 mL of deionized water for 5 min at 80 °C and stirring for 5 min using a hot plate equipped with a magnetic stirrer. The extract was filtered through Whatman No. 1 filter paper and cotton to remove particulate matter, and the brown solution was stored at 4 °C for further use. It remained usable for 1 week, after which gradual loss of plant extract viability occurred [12].

**Figure 1:**
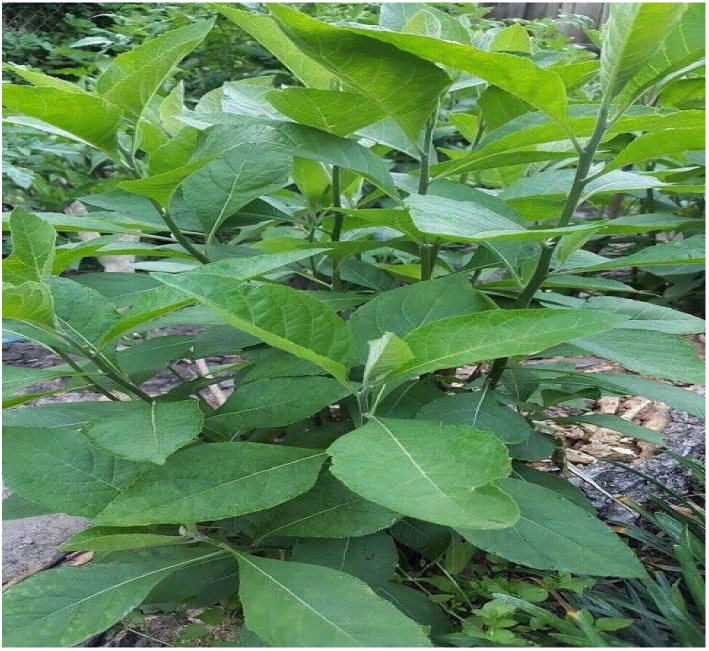
Vernonia amygdalina

### Synthesis of Silver Nanoparticles

Silver nitrate (AgNO_3_) was obtained from Sigma-Aldrich (Germany). Deionized water was used for all reactions. All glassware was washed with dilute nitric acid (HNO_3_) and deionized water and dried in a warm air oven. Different experiments were carried out for three extract quantities (5, 10, and 15 mL), using 50 mL of different concentrations of AgNO_3_ (10^-3^ M, 10^-2^ M, and 10^-1^ M) at room temperature and pressure. The resulting solutions were hand shaken for 1 min, wrapped with aluminum foil, and incubated in the dark to minimize the photoactivation of silver ions. The reactions were performed under static conditions. The color of the solutions changed from initial light brown (which is the color of the *V. amygdalina* leaf extract) to orange in 5 minutes. The first hour of the reaction was monitored by UV-Vis spectroscopy of 2.5 mL of the reaction suspension using a UV-Vis Uviline 9100 spectrophotometer operating at 1 nm resolution, during 300 s with an optical length of 10 mm. After 24 h of incubation, the *Vernonia amygdalina* leaf extract-mediated silver nanoparticles (VA-AgNPs) concentrations were determined by centrifugation. PXRD was carried out using a Rigaku MiniflexII instrument by preparing a thin film of the silver-organic nanopowder on a silicon substrate.

### Determination of the Minimum Inhibitory Concentration (MIC) and the Minimum Bactericide Concentration (MBC)

To evaluate the antimicrobial activity, a multidrug-resistant clinical isolate *of E. coli* was tested against suspensions of VA-AgNPs. This method assesses the sensitivity of bacteria and yeasts to different samples according to the microdilution method described in the ISO standard ISO 20776-1, 2019, and amended by the Antibiotic Committee of the French Society of Microbiology [13]. The EUCAST 2019 microdilution method was adapted to assess the antimicrobial activity of the VA-AgNPs. A 1 mL sample (10 mg/mL) was diluted in Muller Hinton Broth, and serial dilutions were performed using a 96-well microplate. Appropriate controls (negative, positive, and sterility) were included. Each well was inoculated with a standardized microbial suspension (approximately 5×10 CFU/mL). Plates were incubated at 35°C for 18–24 h, and the MIC was defined as the lowest concentration with no visible microbial growth [14]. The concentration range of the samples in the reaction medium was 1; 0.5; 0.25; 0.125; 0.0625; 0.0312; 0.0156; 0.0078; 0.0039; 0.0019 mg/mL.

In bacteria, tetrazolium chloride was used as a color indicator, and The MBC was determined by transplanting 50 µL of suspension from each well where no shoots were observed in the corresponding 50µL crop medium. This was the lowest concentration at which no shoots were observed after transplantation.

From these reports, the bacteriostatic or bactericidal nature of a substance was confirmed. When these ratios are greater than or equal to 4, the substance is said to be bacteriostatic; if these ratios are less than 4, the substance is bactericidal. If they are equal to 1, then they are considered absolute bactericidal [15].

### Evaluation of anti-inflammatory activities

#### Animals and ethics

For this study Wistar albinos’ rats of 8 weeks old, weighing 130 - 150 g were housed in a standard polypropylene cage at room temperature (24±2°C) and relative humidity in the light and dark cycle (from 6 am to 6 pm). They were obtained from the rearing facility of the Department of Pharmaceutical Sciences, Faculty of Medicine and Pharmaceutical Science, University of Douala. They were fed with standard pellet food and tap water ad libitum during 1 week of acclimatization. All experiments were carried out according to the protocol approved by the Institutional Ethics Committee of the University of Douala (protocol approval number CEI-UDo/2617/05/2017/T).

#### Carrageenan-induced rat paw edema method

The anti-inflammatory activity of the synthesized VA-AgNPs was evaluated using a carrageenan-induced rat paw edema model, as described by Winter *et al*. [16]. The experimental animals were randomly divided into five groups (n=6). Group I served as a control and received the inflammatory-inducing agent diclofenac. Group II served as the standard (distilled water), and groups III, IV, and V served as the test groups. Acute inflammation was induced by a single subplantar injection of 0.1 mL of carrageenan (1% carrageenan suspended in 0.9% NaCl) into the right hind paw of the rats 1 h after administration (per os) of VA-AgNPs at doses of 100, 200, and 400 μg/kg for the test groups, indomethacin (10 mg/kg) for the standard group, or distilled water (10 mL/kg) for the control group. Diameters were measured using a digital caliper (Hardened Stainless) immediately before and 30 min, 1 h, 2 h, 3 h, 4 h, and 5 h after carrageenan injection. Edema was expressed as an increase in paw volume, and the percentage of inhibition of edema was expressed as a reduction in volume with respect to the control group using Equation 1:

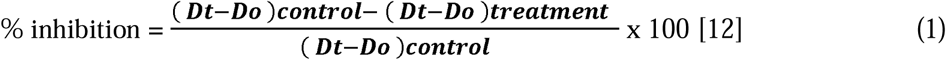

Dt: mean diameter for each group after treatment,

Do: average diameter for each group before treatment.

#### Assessment of acute toxicity on rats of the Wistar strain

The VA-AgNPs toxicity test was performed according to OECD guideline 425. These were limit tests at a dose of 2000 mg/kg in Wistar rats. Batches of three (3) female rats were used for each sample tested, and one batch represented the control group, in which distilled water was administered [17].

The rats were evenly distributed in batches with masses between 120 and 180 g before being subjected to a non-water fast for 12 h prior to sample administration.

The samples and distilled water were administered in one dose by gavage using a gastric tube after calculating the concentrations and volumes to be administered according to formula 2:

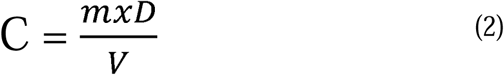

C: Concentration;

D: Dose;

m: Mass of the rat;

V: Volume of solution to be administered. (1 mL for a 100 g rat)

Clinical parameters were monitored at regular time intervals during the hours after gavage, and the resumption of feeding of rats took place 4 h after administration. Simultaneously, the weight of the rats was monitored every day, and their growth rate was calculated using the following formula 3:

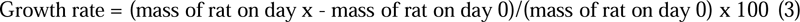

Fourteen (14) days later, the rats were anesthetized with a ketamine solution, and their blood was drawn after incision of the carotid artery for analysis of biochemical parameters. Some organs, such as the liver, kidneys, lungs, heart, and spleen, were removed using surgical scalpel blades, forceps, and scissors.

## RESULTS

### Ultraviolet-Visible Spectroscopy

UV-Visible spectroscopy confirmed the rapid formation of VA-AgNPs, evidenced by a distinct color change from light brown to dark brown upon reaction (Figure 2). Characteristic surface plasmon resonance (SPR) peaks, indicative of Ag nanoparticle synthesis, were observed between 400 and 450 nm (Figure 3). Further analysis revealed that the SPR band intensity, reflecting VA-AgNPs formation and stability, increased significantly with higher pH, increased extract volume, and elevated silver nitrate concentrations (Figure 4, Figure 5).

**Figure 2:**
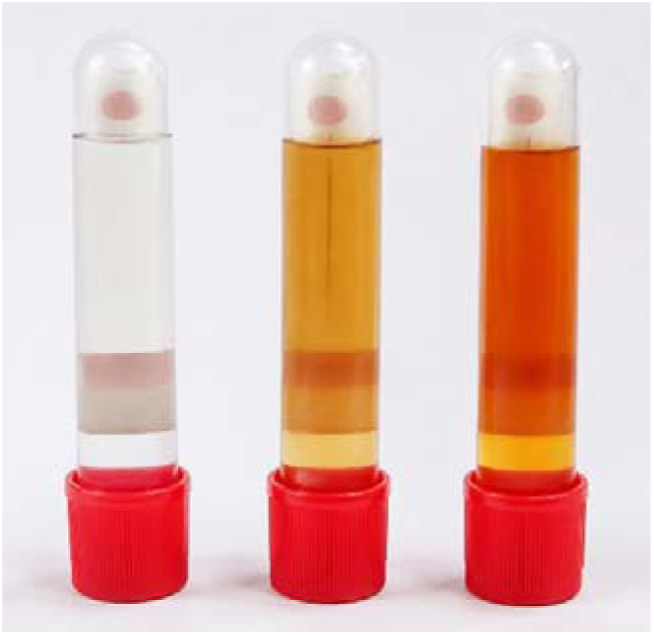
Silver nitrate, *Vernonia amygdalina* leaf extract and VA-AgNPs

**Figure 3:**
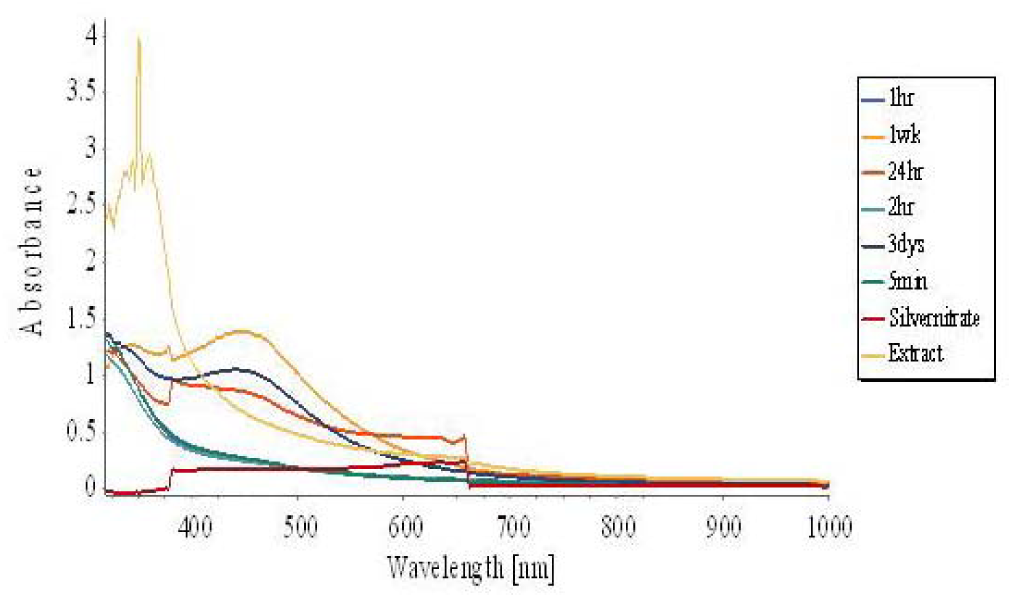
UV-Vis spectra of VA-AgNPs with respect to time

**Figure 4:**
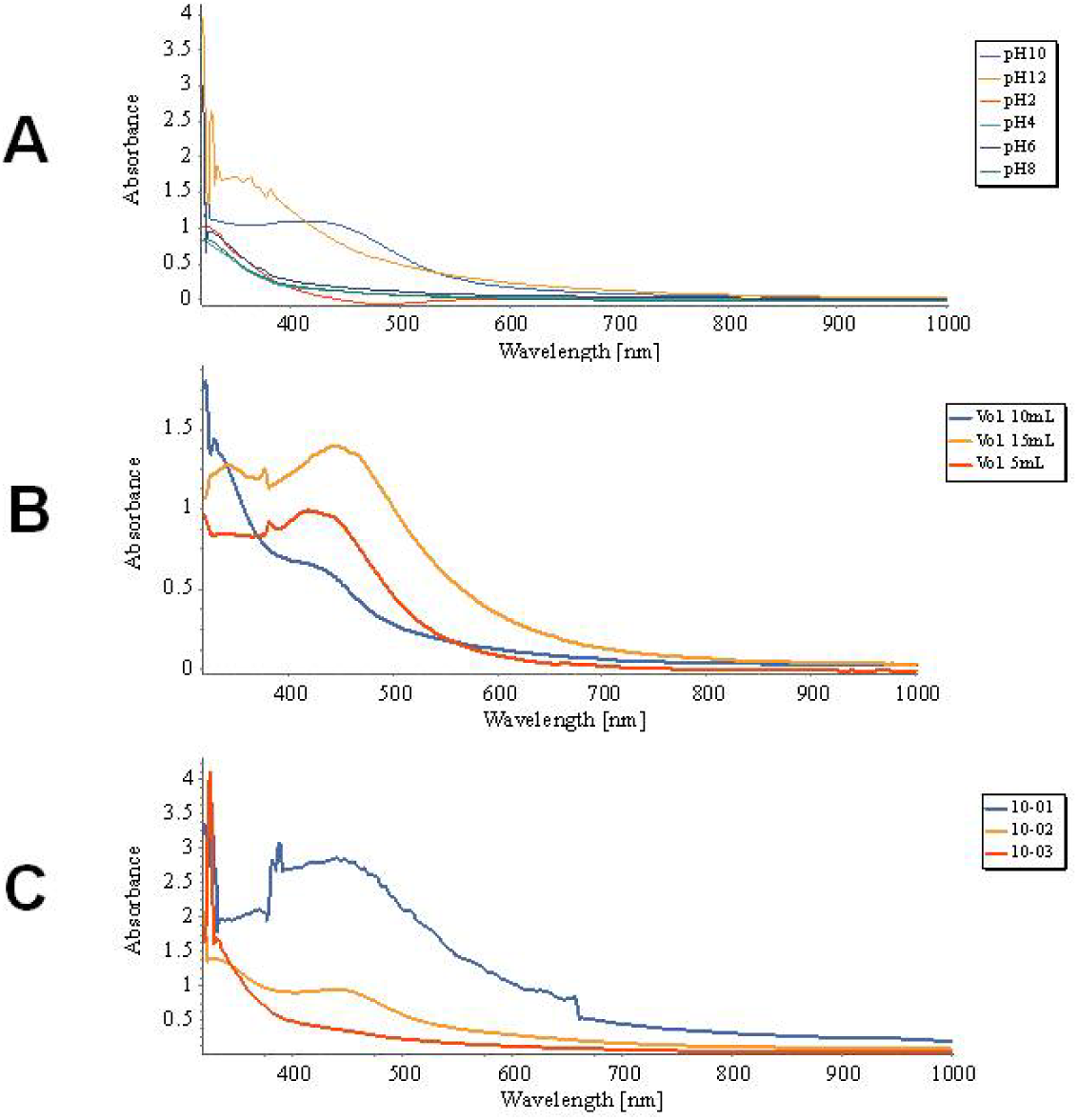
A, B and C Increase in the intensity of the SPR band with an increase in pH, extract volume and silver nitrate concentration, respectively.

**Figure 5:**
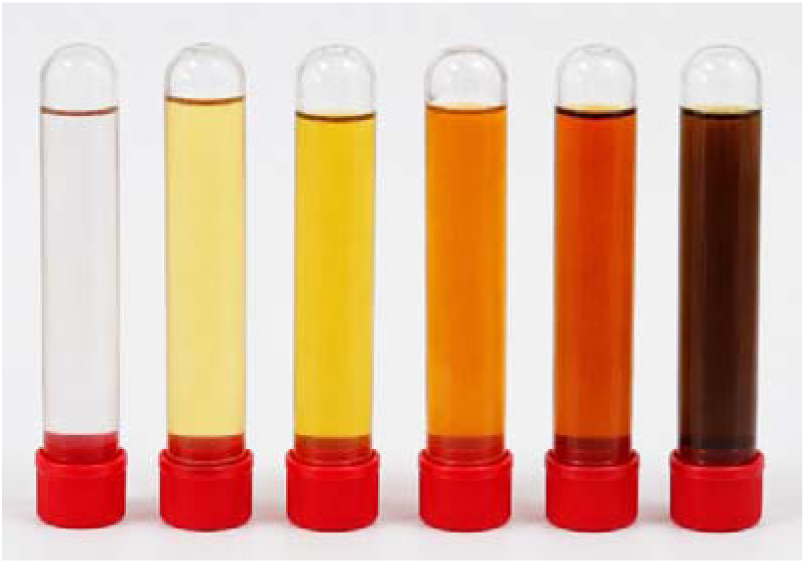
Variation of colors according to pH; pH2, pH4, pH6, pH8, pH10, pH12 (left to right)

### Powder X-ray diffractometry

The powder X-ray diffractogram is presented in Figure 6. It shows 2θ angles ranging from 20 to 60 degrees, appearing a series of crystalline plane with the presence of peaks at 27.7°; 32.1°;38.1°; 44.2° 46.2°; 54.7° and 57.3°.

**Figure 6.**
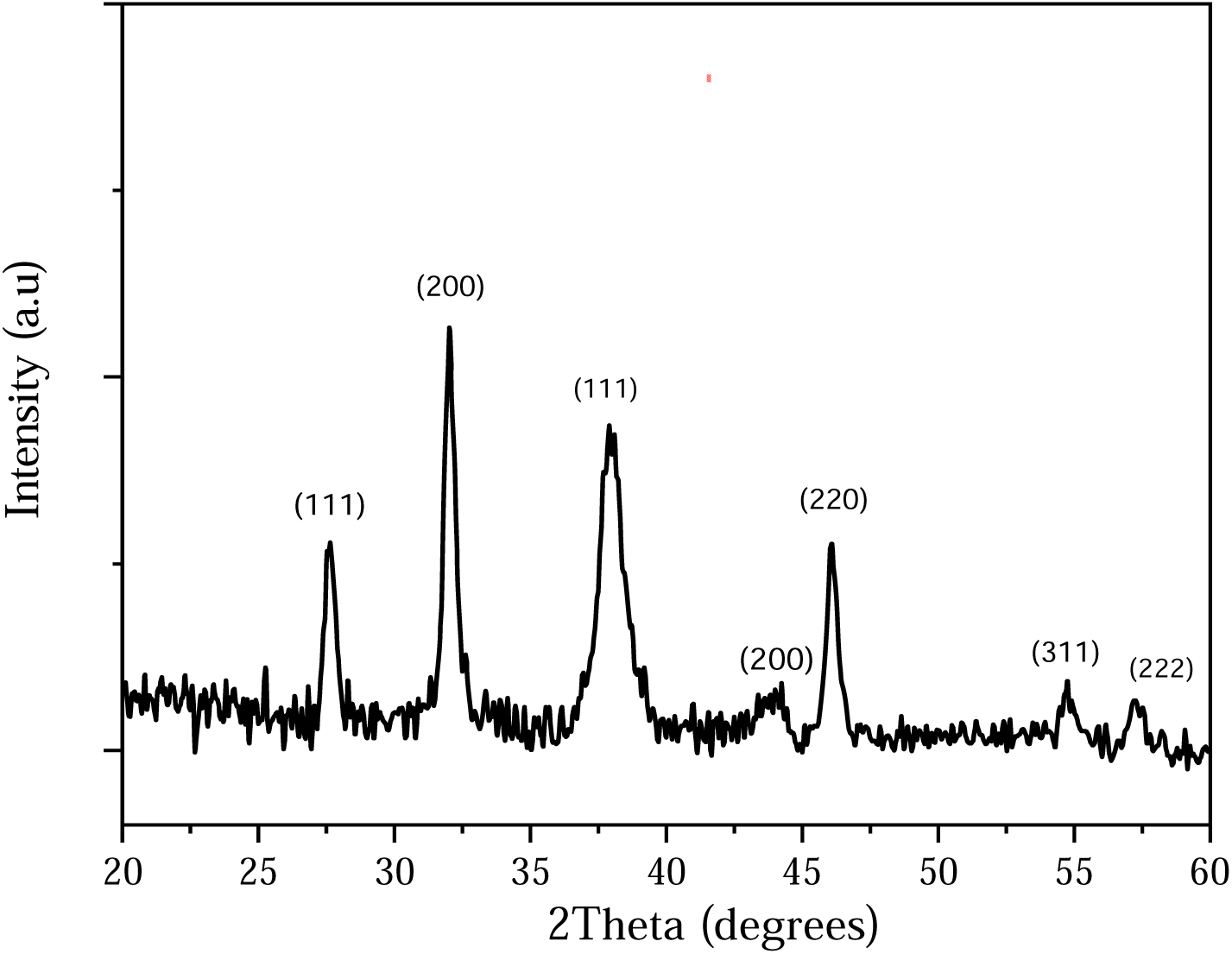
PXRD Spectrum of AgNPs

The average crystallite size of the synthesized NP was determined using the Scherrer equation:

Where D is the crystallite size (nm), λ is the X-ray wavelength (λ = 1.5406 Å), β is the Full width at Half Maximum (FWHM) of the diffraction peaks measured in radians, and θ is the

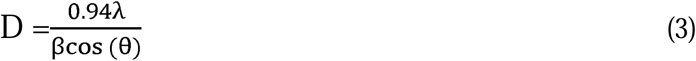

Bragg diffraction angle. No other characteristic peaks were observed in the XRD patterns, indicating the high purity of the as-prepared Ag and AgCl nanoparticles. To calculate the average crystalline particle size of the synthesized Ag and AgCl nanoparticles, we preferred the intense peaks of Ag and AgCl. 2θ values of 38.05° and 44.25°, which can be indexed to the (111) and (200) planes of the face-centered cubic (FCC) structure, respectively (ICDD file: 04-). The XRD pattern also showed the presence of the cubic phase of silver chloride at 2θ values of 27.68°, 32.12°, 46.17°, 54.75°, and 57.30°, corresponding to the planes (111, 200, 220, 311) and (222), respectively (ICDD file: 31-) [11]. The calculated average crystalline particle sizes were 13 and 18 nm for Ag and AgCl, respectively.

### Scanning electron microscopy (SEM)

SEM of VA-AgNPs presents spherical aggregates (Figure 7) with smooth surface morphology.

**Figure 7.**
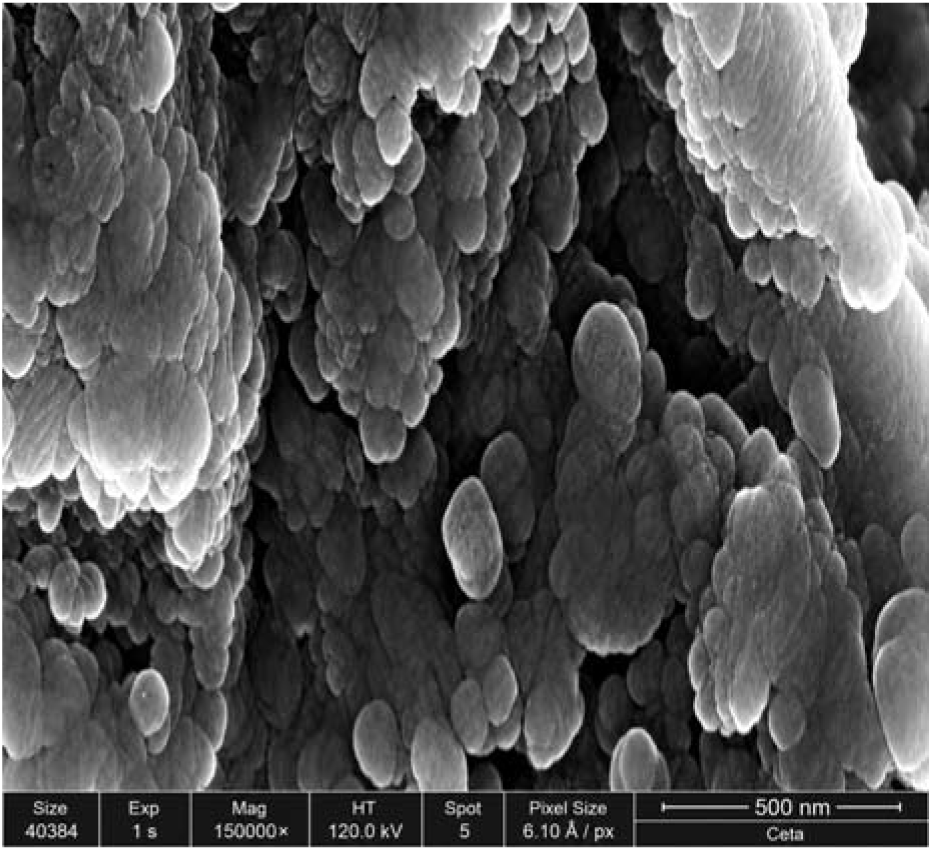
SEM images of AgNPs

### Microbiology control sample

VA-AgNPs and the aqueous extract have been subjected to microbiological control to ensure that they are not contaminated with germs that may grow on culture media used for antimicrobial testing. Table 1 represents observations of culture media impregnated with samples after 48 hours of incubation.

**Table 1.** Microbiological control of samples.

| Samples | Muller Hinton | Sabouraud | Chapman | Hektoen | MBE |
| --- | --- | --- | --- | --- | --- |
| VA-AgNPs (10 mg/mL) | A | A | A | A | A |
| Aqueous Extract | A | A | A | A | A |

A: Lack of shoots

After seeding samples in the Muller Hinton, sabouraud, chapman, hektoen, and MBE mediums, no shoots were observed after 48 hours of incubation.

### The minimum inhibitory and bactericidal concentrations

Evaluation of the antimicrobial activity of the synthesized sample in the multiresistant *E. coli* strain gave values of MIC and MBC, which was compared with the references (ciprofloxacin and aqueous extract). These values are recorded in Table 2.

**Table 2.** MIC and MBC of the sample tested on *E. coli* multiresistant strain compared to ciprofloxacin.

| Microbial strain | *Escherichia coli* |  |  |
| --- | --- | --- | --- |
|  | MIC (mg/mL) | MBC (mg/mL) | MBC/MIC |
| Aqueous Extract (10 mg/mL) | 1,000 | □ 1,000 | / |
| VA-AgNPs (10 mg/mL) | 0,125 ± 0,000 | 0,125 ± 0,000 | 1,000 # |
| Ciprofloxacin (10 mg/mL) (Positive control) | 0,004 ± 0,000 | 0,004 ± 0,000 | 1,000 # |
| Negative control | + |  |  |
| Sterility control medium | - |  |  |

No shoot (-); The presence of a shoot (+); Bactericidal (#)

### Acute oral toxicity

Animals treated with VA-AgNPs survived (Figure 7). This result suggests that the median lethal dose (LD_50_) of VA-AgNPs is higher than the maximum dose administered (2000 mg/kg). According to the OECD, VA-AgNPs are not classified into the category of products at risk of acute toxicity. The curves do not show signs of anorexia; thus, the test models are growing well. The analysis of organ masses shows no differences between the control and test groups (Figure 8). Urea and creatinine do not show statistically different behavior which means that the kidney of treated groups was in good health compared to the control (Figure 9). ASAT and ALAT values allow us to confirm that the liver is not damaged by the nanoparticle treatment (Figure 10).

**Figure 7.**
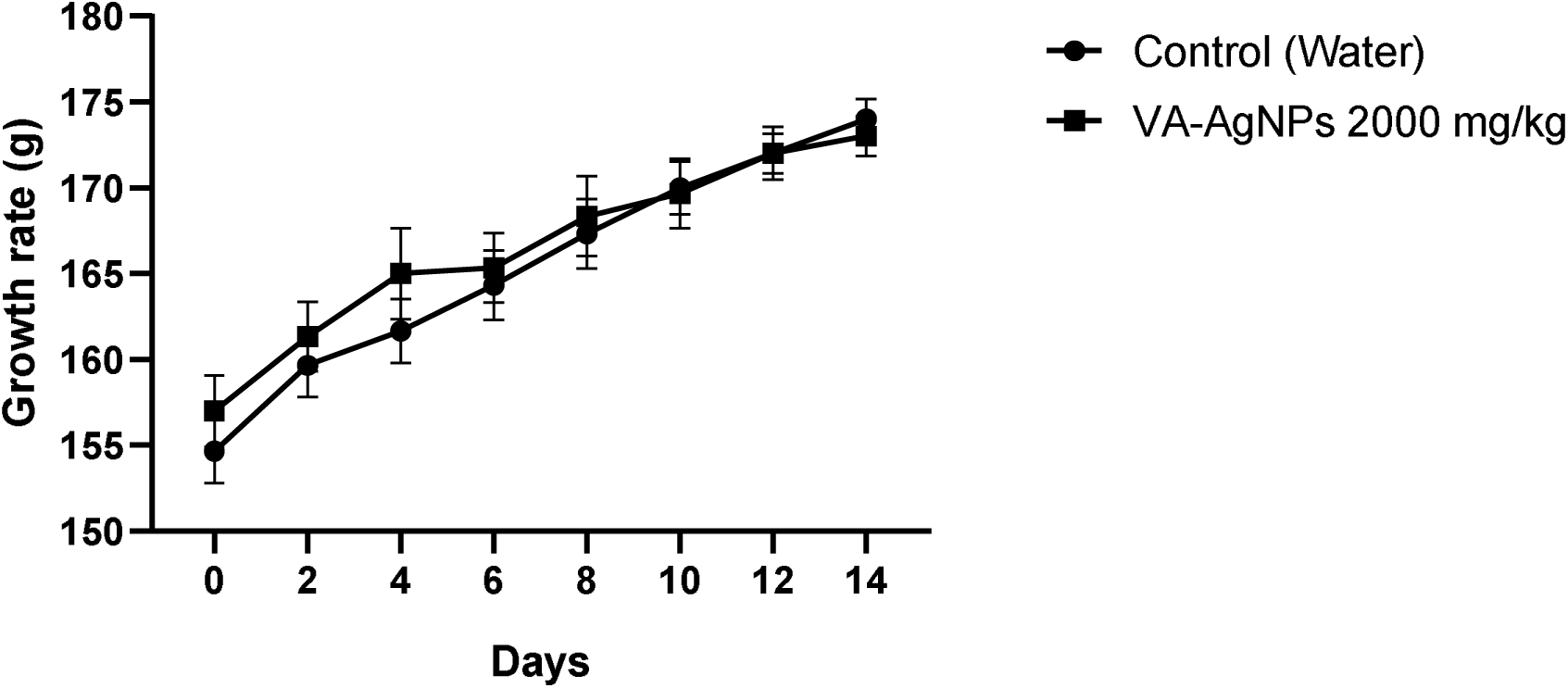
Growth rate of the animals during the acute toxicity assay. The mean of 3 measurement ± standard error of the mean is shown. Two-way ANOVA was used for comparison. The evolution of weight reveals a normal increase in rat weight, without significant differences between the control and test batches at each day.

**Figure 8.**
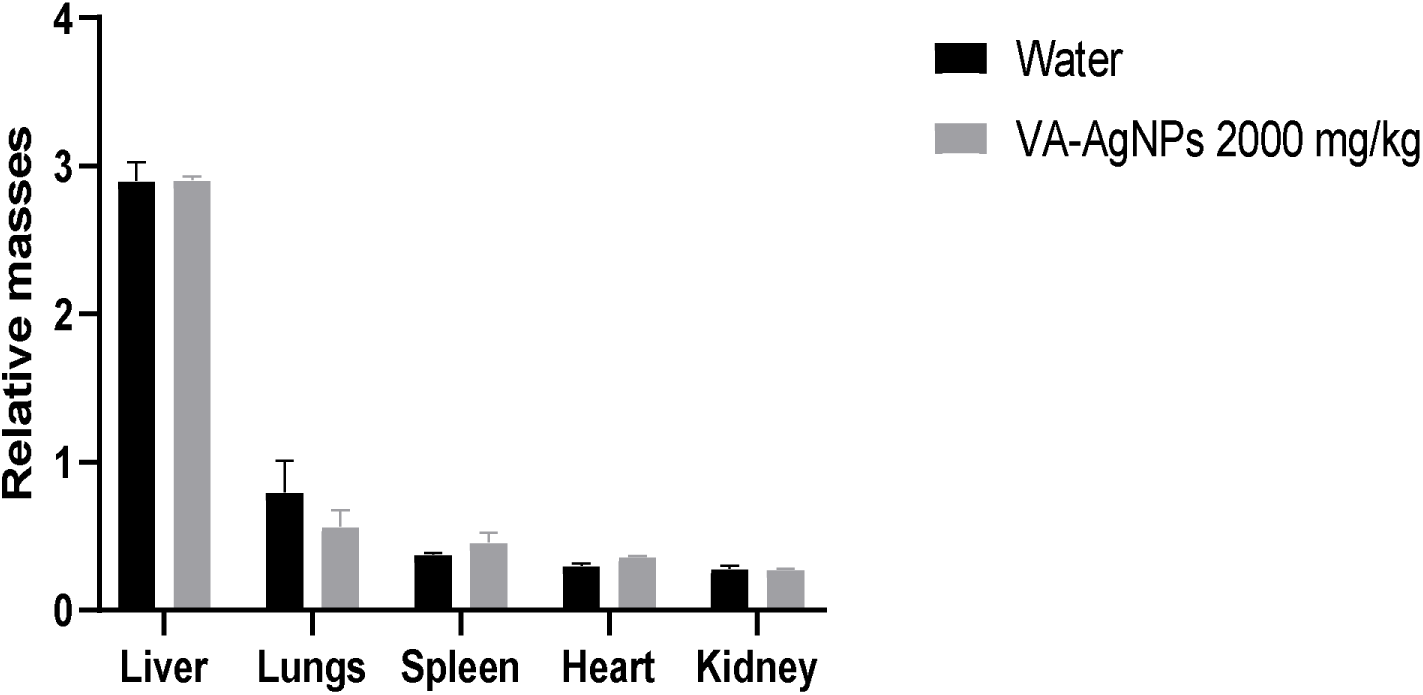
Comparison of the relative masses of the organs of rats tested with VA-AgNPs compared to the control batch presenting no statistically significant differences between the test and the control group. The mean of 3 measurement ± standard error of the mean is shown.

**Figure 9.**
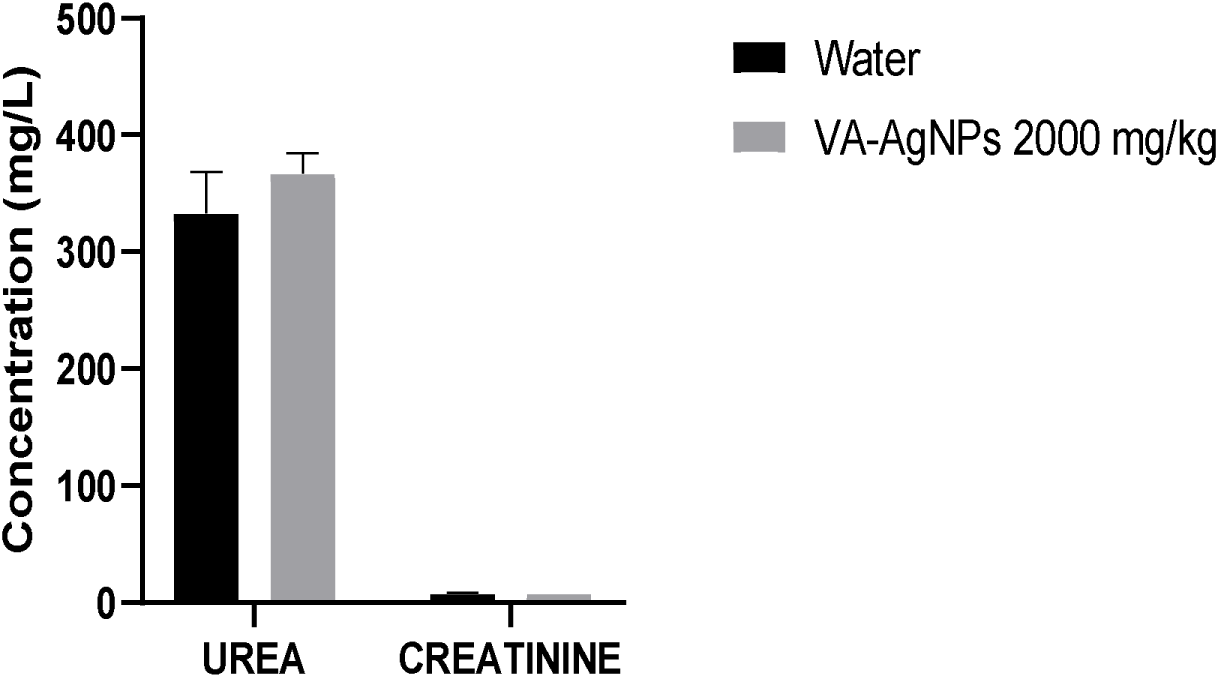
Comparison of creatinemia and uremia of rats subjected to the acute toxicity test for the test batch compared to the control batch without statistically significant differences between the test and the control group. The mean of 3 measurement ± standard error of the mean is shown.

**Figure 10.**
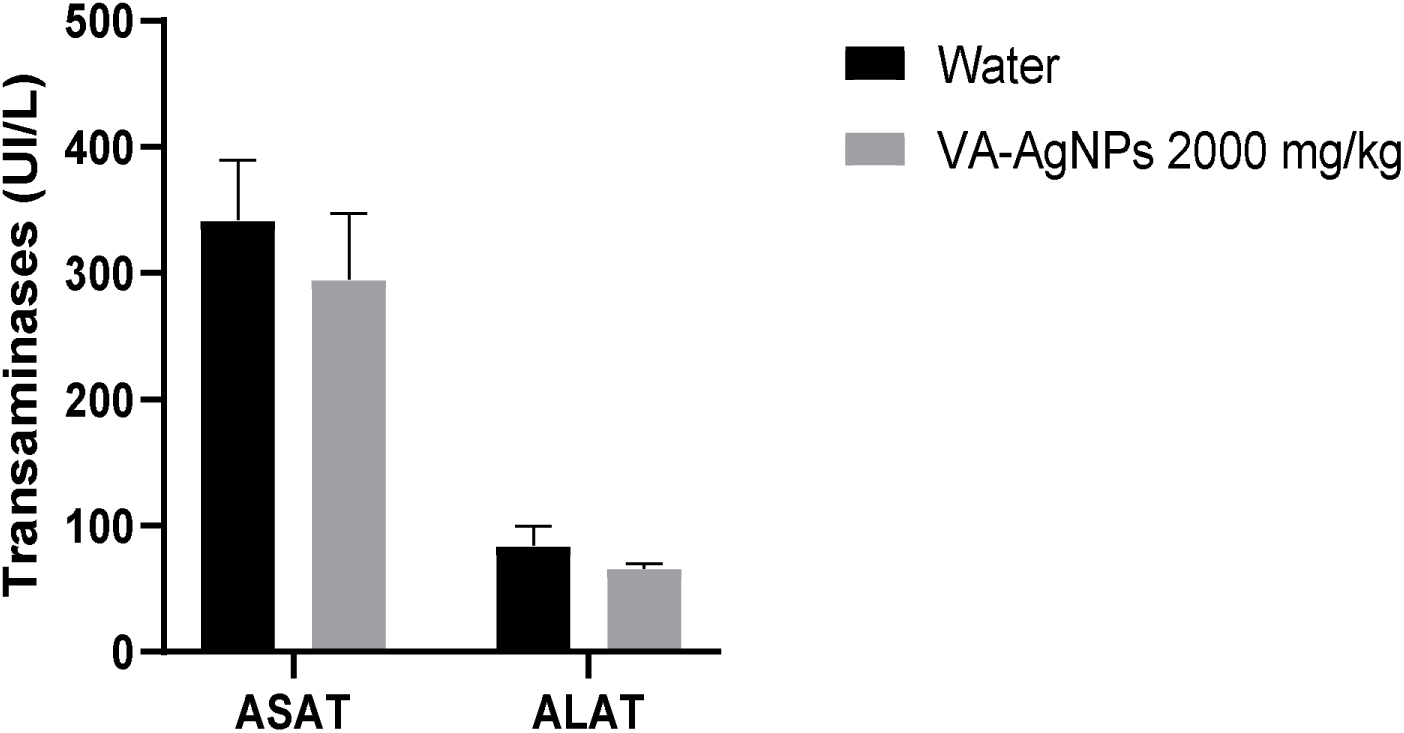
Serum transaminases of rats subjected to the acute toxicity test for the test and the control batch without statistically significant differences between the test and the control group. The mean of 3 measurement ± standard error of the mean is shown.

### Anti-inflammatory activity

The anti-inflammatory activity of VA-AgNPs was evaluated using the carrageenan-induced hind paw edema model in rats, a well-established experimental model of acute inflammation. As shown in Table 3, oral administration of VA-AgNPs produced a marked and dose-dependent reduction in paw edema compared with the untreated inflammatory control group throughout the observation period. Animals treated with VA-AgNPs exhibited significantly lower edema volumes than the control group, indicating effective suppression of the inflammatory response. The highest anti-inflammatory activity was observed at the dose of 400 µg/kg, which produced 96% inhibition of paw edema at 6 h after carrageenan injection. Lower doses also reduced edema formation, although to a lesser extent.

**Table 3.** Effect of VA-AgNPs on rat paw.

| Traitement | Dose (mg/kg) | Diamètre ±SEM (cm) | | | | | | |
| --- | --- | --- | --- | --- | --- | --- | --- | --- |
|  |  | 30 min | 1 h | 2 h | 3 h | 4 h | 5 h | 6 h |
| **Control** | — | 0.9±0.5 | 0.9±0.4 | 0.8±0.4 | 1.0±0.4 | 0.8±0.3 | 0.6±0.3 | 0.7±0.3 |
| **Diclofénac** | 10 | 0.8±0.4<sup>b</sup><br>(5) | 0.7±0.4<sup>b</sup><br>(18) | 0.5±0.4<sup>b</sup><br>(24) | 0.7±0.4<sup>b</sup><br>(27) | 0.3±0.3<sup>b</sup><br>(64) | 0.27±0.00<sup>b</sup><br>(56) | 0.10±0.05<sup>b</sup><br>(86) |
| **VA-AgNPs** | 0.1 | 0.9±0.2<sup>c</sup><br>(-4) | 0.8±0.2<sup>c</sup><br>(7) | 0.7±0.2<sup>c</sup><br>(11) | 0.06±0.00<sup>c</sup><br>(94) | 0.06±0.05<sup>c</sup><br>(93) | 0.12±0.00<sup>c</sup><br>(80) | 0.11±0.00<sup>c</sup><br>(84) |
| **VA-AgNPs** | 0.2 | 1.0±0.2<sup>c</sup><br>(-14) | 0.7±0.3<sup>c</sup><br>(21) | 0.5±0.3<sup>c</sup><br>(27) | 0.13±0.00<sup>c</sup><br>(87) | 0.1±0.0<sup>c</sup><br>(88) | 0.07±0.00<sup>c</sup><br>(89) | 0.05±0.04<sup>c</sup><br>(93) |
| **VA-AgNPs** | 0.4 | 0.8±0.2<sup>c</sup><br>(5) | 0.7±0.3<sup>c</sup><br>(20) | 0.5±0.3<sup>c</sup><br>(31) | 0.3±0.2<sup>c</sup><br>(72) | 0.18±0.00<sup>c</sup><br>(78) | 0.05±0.00<sup>c</sup><br>(92) | 0.03±0.00<sup>c</sup><br>(96) |

b (P < 0.01), c (P < 0.001)

## Discussion

The color change of the plant extract and silver ion solution mixture was first observed through visual monitoring. The solution color changed within seconds to pale yellow and then brown due to the formation of plasmons at the colloid surface, indicating the synthesis of silver nanoparticles. The same sharp surface plasmon resonance absorbance band was obtained with different extract concentrations (5, 10, 15 mL) at 10^-3^ M AgNO_3_. Then, 5 mL of extract was sufficient to completely reduce 50 mL of silver ions at a concentration of 10^-3^ M (Figure 4). Plasmon resonance absorbance increases when a 10^-2^ M AgNO_3_ solution is used with different extract concentrations, then more silver ion is available for reduction. At 10^-1^ M concentration of silver ions with different *V. amygdalina* extract concentrations, the nanoparticles aggregate because of the deficiency of leaf extract molecules to act as capping agents. The barrier potential developed because of the competition between the weak van der Waals forces of attraction and electrostatic repulsion is broken [12]. As postulated by Mie’s theory, spherical nanoparticles result in a single surface plasmon resonance (SPR) band in their absorption spectra. In contrast, anisotropic particles provide two or more SPR bands, depending on their shape [15]. In the present study, the reaction mixtures confirmed single SPR bands, revealing the spherical shape of VA-AgNPs, which tended to become anisotropic with time. The plant leaf extract of *V. amygdalina* acted as a reductant and stabilizer for the silver nanoparticles. The sharp surface Plasmon resonance band increased with the silver ion concentration, as observed for the *Megaphrynium macrostarchyum* leaf extract [18].

Time-recorded spectra between 5 min and 1 week for a mixture of 3 mL of the aqueous extract and 10 mL of 1 mM silver nitrate solution showed a wide range of VA-AgNPs SPR peaks between 350 and 550 nm. The absorption intensity of the SPR band increased over time and started forming a plateau from 24 h with an absorption intensity of 0.9 until it reached its highest plateau in one week with an absorption intensity of 1.5. The maximum absorbance of VA-AgNPs using *V. amygdalina* was in the range of 400-450 nm due to the surface plasmon resonance [19]. The UV-Vis spectra also revealed that VA-AgNPs were formed rapidly within a few minutes, indicating that *V. amygdalina* accelerated the biosynthesis of silver nanoparticles. This rapid reduction was observed in previous research for carob leaf extract within 2 min [20].

The reaction between Ag^+^ and the reducing material in the extract was followed for one week, and UV-Vis spectra of VA-AgNPs were recorded as a function of time after the addition of different amounts of *V. amygdalina* leaf extract. The reduction, as well as the nucleation and growth of the nanoparticle size, increased from 24 h to 1 week, but polydispersions occurred. The reaction time resulted in a gradual increase in the absorbance bands. The color intensity of the solution changed from light yellow to deep brown at the end of the reaction because of the increasing amount of VA-AgNPs and aggregation [18]. The absorbance increased with pH, which can be attributed to the increase in the production of colloidal silver nanoparticles and the reduction rate. Absorbance did not decrease at pH values higher than 8, as observed for *Megaphrynium macrostachyum* leaf extract [18]. Furthermore, the brown color of the nanoparticles appeared shortly after mixing AgNO_3_ with the extract at pH 4-12. As observed in the bark extract of *Pinus eldaria,* pH affects the amount of nanoparticle production and its stability and is a critical factor in controlling the size and morphology of nanoparticles [21]. Again, the pH influenced the rate of reduction. The reaction mixture turned brown when silver was reduced, and the color of the reaction mixture accelerated with increasing pH. At pH 10, the sharp surface plasmon resonance band indicated the occurrence of a monodisperse suspension. Previous investigations have shown that the size and shape of biosynthesized nanoparticles can be manipulated by varying the pH of the reaction mixtures [18]. A major influence of the pH of the reaction is its ability to change the electrical charges of biomolecules, which could affect their capping and stabilizing abilities, and then the growth of the nanoparticles [22]. The high pH environment eased the reduction and stabilization ability of antioxidants in the *V. amygdalina* leaf extract, a situation also found in olive leaf extract. The number of nuclei increased with increasing pH due to the promoted reactivity of the *V. amygdalina* leaf extract reductants.

X-ray diffractometry was performed on VA-AgNPs. It revealed the nanocrystalline structure of the formed nanoparticles composed of Ag and AgCl with 13 and 18 nm grain sizes respectively. The SEM image shows a spherical shape for the sample with a cluster of aggregation.

Beyond their structural and optical properties, the synthesized VA-AgNPs were subsequently evaluated for their crucial biological activities, including antibacterial efficacy, acute toxicity, and anti-inflammatory potential.

The resistance of human pathogens to the commercially available antimicrobial agents and antibiotics has raised the need to explore new natural and inorganic substitutes to overcome the problem [23]. More et al. [24] demonstrated that Ag nanoparticles are attached to the cell surface of *E. coli* and then penetrate the cell, destroy the cell cytoplasm, and kill the organism. Ag nanoparticles significantly increase cell permeability and affect proper transport through the plasma membrane [24]. It appears that sulfur and phosphorus from proteins or enzymes or phosphorus moiety of DNA of the bacterial system may be affected by VA-AgNPs that lead to the inhibition of the enzyme system of the organism [24].

While the VA-AgNPs demonstrated a bactericidal effect, their MIC was substantially higher than that of ciprofloxacin against the multiresistant *E. coli* strain (Table 2), indicating a lower potency compared to the reference antibiotic. This suggests that while promising, further optimization may be needed to achieve comparable efficacy. These activities are related to the possibility of releasing the silver ions directly into the microbial cell that will be able to act on various sites [24]. Synthesized nanoparticles are covered with molecules from the extract that may also have intrinsic antimicrobial properties. Therefore, VA-AgNPs would also play a good role as a vector for delivering Ag^+^ ions and molecules from the extract at their site of action.

Concerning acute oral toxicity, all animals treated with VA-AgNPs survived (Figure 7). This result suggests that the median lethal dose (LD_50_) of VA-AgNPs is higher than the maximum dose administered (2000 mg/kg). According to the OECD, VA-AgNPs are not classified into the category of products at risk of acute toxicity. This result corroborates the results of Tchangou Njiemou et al. (2020) who showed that nanoparticles are not toxic at a dose of 2000 mg/kg [7]. The curves do not show signs of anorexia; thus, the test models are growing well. The analysis of organ masses shows no differences between the control and test groups (Figure 8). Also, urea and creatinine do not show statistically different behavior which means that the kidney of treated groups was in good health when compared to the control (Figure 9). ASAT and ALAT values allow us to confirm that the liver is not damaged by the nanoparticle treatment (Figure 10).

Carrageenan-induced edema is a triphasic response that involves the release of different mediators, including histamine and serotonin, in the first phase (1 hour). The second phase is mediated by the release of kinins (2 hours), and the third phase is attributed to prostaglandins and cyclooxygenase products, which last for 3 to 5 hours [25, 26]. These chemicals increase vascular permeability during their production, thus promoting the accumulation of fluid in tissues that explains edema [26]. Oral pretreatment of animals with VA-AgNPs resulted in effective inhibition of the edema rate in all phases compared to the control (Table 3). Maximum inhibition (96%) was obtained at a dose of 400µg/kg, 6 hours after injection. This result suggests that synthesized nanoparticles may act by inhibiting the release of histamine, serotonin, and kinins and/or by interfering with the synthesis of prostaglandin mediators as previously reported by Belle Ebanda Kedi *et al*. [12]. This result is different from that of Adiukwu *et al*. in 2013 who demonstrated the anti-inflammatory and antipyreticactivity of the leaf, root and saponin fraction of *V. amygdalina* and who had a percentage inhibition of edema of 88% at 800 mg/kg of the aqueous extract of *V. amygdalina* [27]. Despite the promising findings, this study has several limitations. The antibacterial activity was evaluated against a single strain of *E. coli*; future work should include a broader spectrum of bacterial pathogens and clinical isolates to confirm broad-spectrum efficacy. While acute toxicity was assessed, long-term toxicity and pharmacokinetics studies are essential to fully understand the safety profile of VA-AgNPs for potential therapeutic applications. The precise mechanism of action for both antibacterial and anti-inflammatory effects was not fully elucidated, necessitating further investigations into cellular and molecular pathways.

## CONCLUSION

This study demonstrates the successful green synthesis of silver nanoparticles (AgNPs) using *Vernonia amygdalina* leaf extract as both reducing and stabilizing agent. The rapid formation of stable, spherical *Vernonia amygdalina* leaf extract-mediated silver nanoparticles (VA-AgNPs) was confirmed by UV-Vis spectroscopy, with a sharp surface plasmon resonance band appearing within minutes, and intensifying over time, extract concentration, silver ion concentration, and pH. X-ray diffraction analysis confirmed the crystalline nature of the nanoparticles, with average crystallite sizes of 13 and 18 nm, for Ag and AgCl respectively. SEM analysis further revealed spherical morphology with moderate aggregation.

Functionally, the biosynthesized AgNPs displayed strong antibacterial activity against multidrug-resistant *E. coli*, with a MIC and MBC of 0.125 mg/mL, indicating a potent bactericidal effect. In vivo toxicity evaluation revealed no adverse effects at 2000 mg/kg, indicating a favorable safety profile. Notably, the VA-AgNPs exhibited significant anti-inflammatory activity in a carrageenan-induced rat paw edema model, achieving up to 96% inhibition at 400 µg/kg, superior to both the aqueous extract and previous reports using *V. amygdalina* alone.

These findings highlight the potential of *V. amygdalina*-derived VA-AgNPs as a promising, eco-friendly nanomaterial for antimicrobial and anti-inflammatory applications, with high biocompatibility and therapeutic efficacy. Future investigations should explore their mechanisms of action, long-term safety, and potential for formulation into clinical products.

## Declarations

### Data availability

The authors declare that the data supporting the findings of this study including raw data files are available from the corresponding author upon reasonable request.

### Statement on animal rights

On behalf of all authors, the corresponding author affirms that animal rights were upheld in the study. Ethics approval Ethical clearance Nr. CEI-UDo/2617/05/2017/T was obtained from the Institutional Research Ethics Committee for Human Health of the University of Douala (CEI-UDo).

